# Defective DcpS Decapping Manifests in Creatine Deficiency Syndrome and Neurological impairment

**DOI:** 10.1101/2024.12.18.629192

**Authors:** Jun Yang, Geeta Palsule, Xinfu Jiao, Ronald P. Hart, Megerditch Kiledjian

## Abstract

Biallelic mutations in the *DCPS* gene disrupting the decapping activity of the scavenger decapping protein DcpS, leads to neurodevelopmental deficiencies and intellectual disability. However, the molecular basis for the neurogenesis defects in these individuals remains unknown. Here we show that cells derived from individuals with a *DCPS* mutation harbor a creatine deficiency and a corresponding elevation of the creatine precursor, guanidinoacetate (GAA). The altered metabolite levels are a consequence of a reduction in both the mRNA and protein levels for the enzyme that converts GAA into creatine, guanidinoacetate methyltransferase. Importantly, the compromised neurogenesis and neurite outgrowth phenotypes observed during the differentiation of DcpS mutant patient derived induced pluripotent stem cell differentiation into neurons was reversed upon supplementation of creatine monohydrate. These findings suggest creatine deficiency as the major underlying factor for the neurogenetic defect detected in DcpS mutant cells and a potential driver of the neurological deficiencies in affected individuals.

## Introduction

Alteration of gene expression at the level of mRNA turnover is a critical modulator of neural development. A key contributor to the stability and accumulation of an mRNA is the protective N7-methylguanosine (m^7^G) cap at the 5′ end and its removal termed decapping. mRNA decapping is primarily carried out by proteins in three distinct protein families. One is constituted by the Nudix family of enzymes where the Dcp2, Nudt16, Nudt3 and Nudt2 have been demonstrated to function as decapping proteins in cells ^1–5^ while Nudt12, Nudt15, Nudt17 and Nudt19 are known to at least function *in vitro* ^6^. Interestingly, several of the Nudix hydrolases also decap noncanonical caps consisting of NAD caps (Nudt12 and Nudt16) and FAD caps (Nudt2 and Nudt16) ^7^. A second family is the Dxo/Rai1 family of decapping enzymes that function in 5′ end quality control and hydrolyze both canonical and non-canonical capped RNAs ^7^. A third family consists of the Histidine triad (HIT) family of proteins primarily consisting of the DcpS scavenger decapping enzyme ^8,9^. Members of the HIT family can hydrolyze the residual cap structure dinucleotide following mRNA 3′ end decay ^8,10^ and the mRNA cap on an intact RNA ^11^.

In addition to the well-established role of RNA binding proteins in neural function ^12^, the specific significance of decapping enzymes in neurogenesis became apparent with the implication of the Dcp2 decapping stimulatory protein, Edc3 in cognitive disability ^13^ and the causal link between mutations in the *DCPS* gene encoding the DcpS protein in neurodevelopmental and cognitive disorders that could also include microcephaly, musculoskeletal and craniofacial abnormalities collectively referred to as Al-Raqad syndrome ^13–16^. More recent analyses revealed a role for *DCPS* in human neuronal and mouse neocortical development ^17^ . Induced neurons from induced pluripotent stem cells (iPSCs) derived from individuals with compromised DcpS decapping demonstrated curtailed differentiation and neurite projections. Altered DcpS expression in developing mouse brain impacts axonal growth and neocortical subtype identity. The role of DcpS decapping in neural development and neurogenesis underscores its contribution to cognition, yet the underlying mechanism remains unknown.

Creatine (Cr) (α-N-methylguanidino acetic acid), a nitrogenous organic compound along with its phosphorylated derivative, Phospho-Cr, serve as an essential energy currency to transport high energy phosphates and maintain ATP levels ^18,19^. Neuronal cells are among the most energy demanding cells in an organism and consume up to 20% of total energy ^20^. Cr is an important component that enables rapid resynthesis of neural ATP especially during increased metabolic activity ^21^. Consequently, Cr deficiency syndromes primarily manifest in the central nervous system (CNS) and among other consequences, promote cognitive impairment and delays in speech acquisition ^22^, two phenotypes observed in individuals with impaired DcpS decapping activity ^13^.

Here we report cells compromised in DcpS decapping activity have reduced Cr and Phospho-Cr levels. Moreover, supplementation of DcpS mutant patient cells with Cr reversed the decreased neuronal differentiation and neurite elongation, indicating the modulation of Cr may underly the neurogenesis defect in DcpS mutant cells and manifest the cognitive defects in affected individuals.

## Results

### Differential gene expression analysis reveals disrupted creatine metabolism and cell-cell signaling in DcpS mutant cells

A homozygous splice site variant within the *DCPS* gene (c.636+1G>A) disrupts the normal splicing pattern of intron 4, leading to the utilization of a cryptic splice site 45 nucleotides downstream. Translation of the resulting aberrant mRNA produces a mutant variant protein with a 15 amino acid insertion that manifests in a neurodevelopmental disorder ^13^. To investigate the impact of this mutation on gene expression, RNA sequencing (RNA-seq) was performed on lymphocytes derived from patients carrying this mutation. Comparative analysis between patient derived DcpS mutant (DcpS-Mut) cells (individual IV-3 in ^13^) and control cells (DcpS-Ctrl) from a phenotypically normal consanguineous cousin heterozygous for the G>A mutation (individual III-3 in ^13^), revealed significant alterations in the expression profiles of various transcripts. Volcano plots of reference transcripts with at least 2-fold changes in gene expression levels and Σ5% FDR reveal the extent of differential transcript expression between DcpS-Mut and DcpS-Ctrl cells (Figure 1A) with 1160 unique transcripts elevated and 1226 unique transcripts reduced in DcpS-Mut cells.

**Figure 1.**
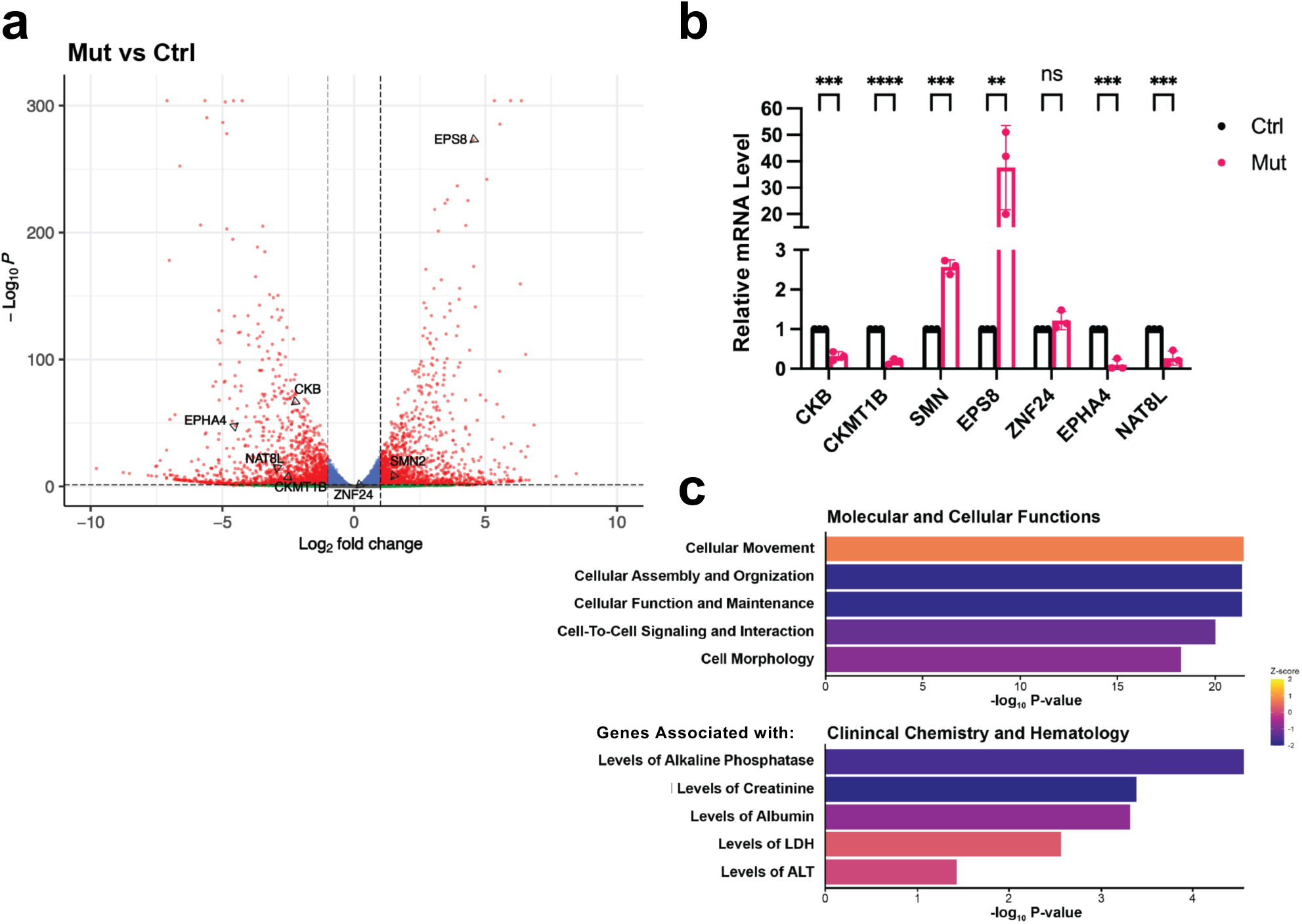
DcpS dependent differential gene expression. **(a)** Volcano plot illustrating the differential gene expression profile identified through RNA sequencing analysis in the indicated lymphoblast cell lines with transcripts tested in panel B indicated. **(b)** Quantification of gene expression via RT-qPCR, with normalization to GAPDH expression levels. Statistical analysis conducted on data from n=3 biological replicates, each with three technical replicates. Error bars represent the Standard Error of the Mean (SEM). Significance levels indicated by asterisks: ****P < 0.0001, ***P < 0.001, **P < 0.01, and ns denoting non- significant differences (P > 0.05), determined by unpaired t-test. **(c)** Functional enrichment analysis of gene categories based on RNA sequencing data plotted with p values on a log10 scale in the X-axis, analyzed using Ingenuity Pathways Analysis (IPA) software.

Validation of the RNA-seq analysis was carried out to quantitate a random subset of transcripts that either increased or decreased directly by quantitative reverse transcript PCR (qRT-PCR). All four mRNAs that decreased (CKB, CKMT1B, EPHA4 and NAT8L) and two mRNAs (SMN and EPS8) that increased by at least two-fold or did not change (ZNF24) in the DcpS-Mut cells by RNA-seq were further confirmed by qRT-PCR (Figure 1B). Functional enrichment analysis using Ingenuity Pathways Analysis of the differentially regulated transcripts highlighted several key pathways, notably cellular signaling, cell-to-cell communication, and amino acid metabolism (Figure 1C). Among these pathways, a reduction of transcripts that would result in the elevation of Cr byproduct, creatinine, was also noted. Interestingly, two of the down regulated transcripts validated, CKB and CKMT1B in Figure 1B, function within the creatine (Cr) biosynthesis pathway. These findings implicated alterations in Cr metabolism as a prominent category of interest.

### Identification of metabolic anomalies in the creatine biogenesis

To determine whether there are metabolomic anomalies in DcpS-Mut lymphoblast cells that may provide potential links to neurological deficiencies, we conducted a comprehensive quantitative metabolomics analysis. The results revealed a marked reduction in intracellular Cr and Phospho-Cr levels in patient-derived cells harboring the DcpS mutation (Figure 2A). Cr is synthesized primarily in a two-step process (Figure 2B). First, arginine and glycine are converted to guanidinoacetate (GAA) by the enzyme L-arginine:glycine-amidinotransferase. GAA is subsequently methylated by guanidinoacetate methyltransferase (GAMT) to produce Cr. Importantly, the metabolomic analysis also revealed a concomitant significant increase in the levels of the creatin substrate, GAA (Figure 2A). These findings suggest that DcpS-Mut cells harbor a disruption in the Cr synthesis pathway and implicate a defect in the transition of GAA to Cr by GAMT.

**Figure 2:**
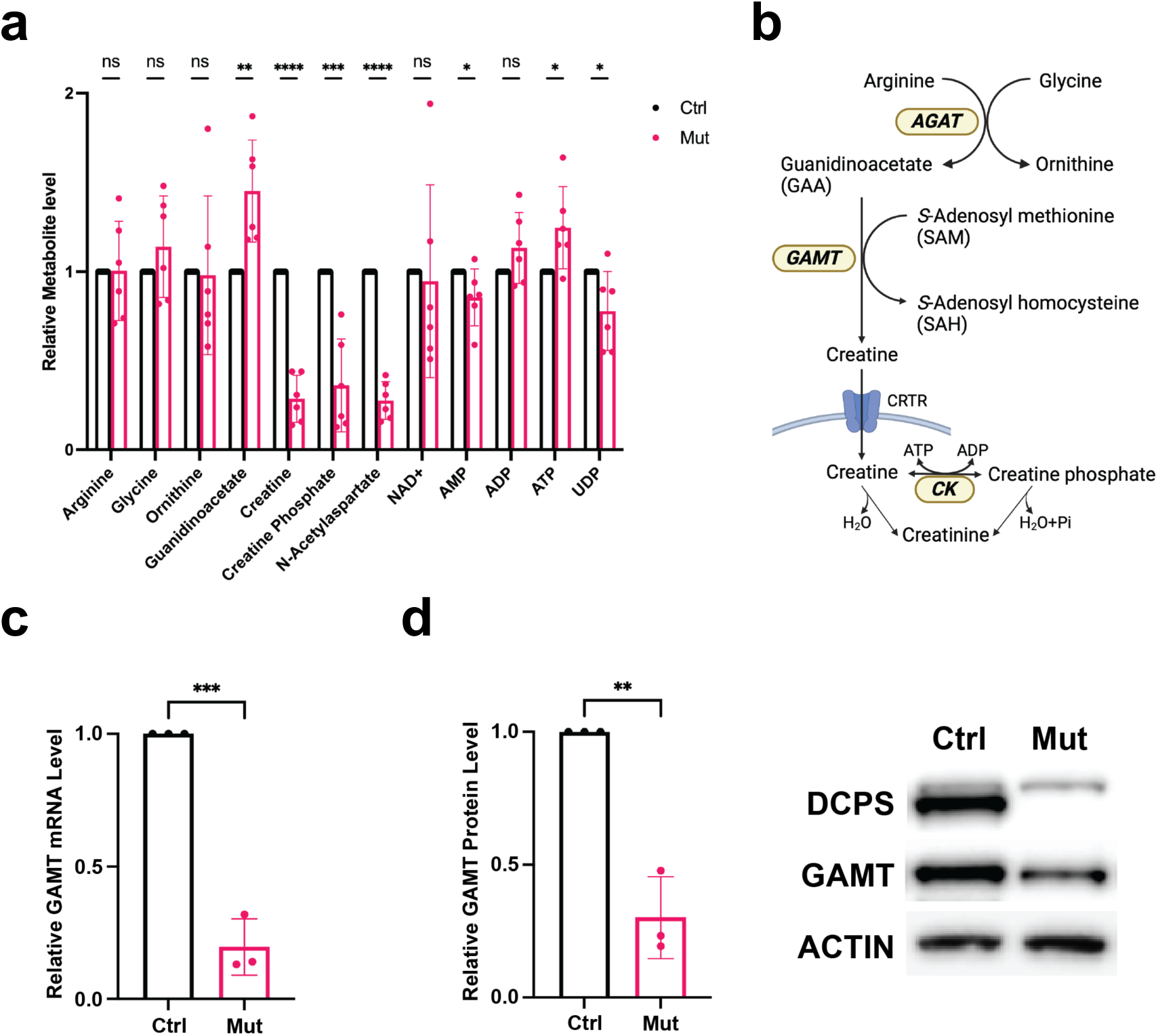
DcpS Mutation Impairs Human Neuronal Development In Vitro, with Restoration by Cr Supplementation. **(a)** Metabolite differences between DcpS-Ctrl versus DcpS-Mut cells. The average of 6 independent metabolic panels are shown. Significance levels are represented by asterisks: ****P < 0.0001, ***P < 0.001, **P < 0.01, *P < 0.05, and ns denoting non-significant differences (P > 0.05). **(b)** Schematic representation of the Cr pathway. **(c)** Quantification of guanidinoacetate methyltransferase (GAMT) mRNA expression by RT- qPCR in DcpS-Ctrl and DcpS-Mut, with normalization to GAPDH expression. Statistical analysis conducted on data from n = 3 biological replicates, each with three technical replicates, using an unpaired t-test. Significance represented by ***P < 0.001. **(D)** Quantification of GAMT protein levels by western blot in DcpS-Ctrl and DcpS-Mut cells normalized to Actin expression. Statistical analysis performed on data from n = 3 biological replicates, with three technical replicates for each, using an unpaired t-test. Significance indicated by **P < 0.01.

To address whether DcpS-Mut cells possess anomalous expression of GAMT, levels of GAMT mRNA and protein were analyzed. Consistent with a disruption in GAMT expression, it’s mRNA was indeed reduced in the RNA-seq analysis and was further validated by direct qRT-PCR (Figure 2C) in the DcpS-Mut cells relative to the control cells. Similarly, GAMT protein levels were concomitantly reduced (Figure 2D). Collectively, our data suggest DcpS-Mut cells harbor an aberrant Cr biosynthesis pathway. Moreover, the anomaly appears to reside primarily through the downregulation of GAMT mRNA and protein levels, leading to an accumulation of guanidinoacetate and a subsequent reduction in intracellular Cr and Phospho-Cr levels in patient-derived cells.

### Creatine addback restores neuronal differentiation in DcpS mutant induced neurons

Induced pluripotent stem cells (iPSCs) derived from DcpS-Mut cells are impaired in their differentiation into induced neuron (iN) cells and project neurite outgrowths that are shorter than control cells ^17^. Given that DcpS-Mut cells contain reduced Cr levels, we sought to determine whether the neurogenesis defects in these cells was a manifestation of the reduced Cr. Using a titration of exogenous Cr supplemented into the culture medium, 100µM of Cr monohydrate was established as optimal to restore DcpS-Mut cell intracellular Cr to levels comparable to that of DcpS-Ctrl cells (Figure 3A). Importantly, the reduction in GAMT mRNA and protein levels observed in Figures 2C and 2D were unaltered upon Cr supplementation (Figures 3B and 3C) indicating any differences potentially observed upon addition of Cr to the DcpS-Mut cells would be a function of the supplemented metabolite and not changes arising from altered GAMT expression.

**Figure 3:**
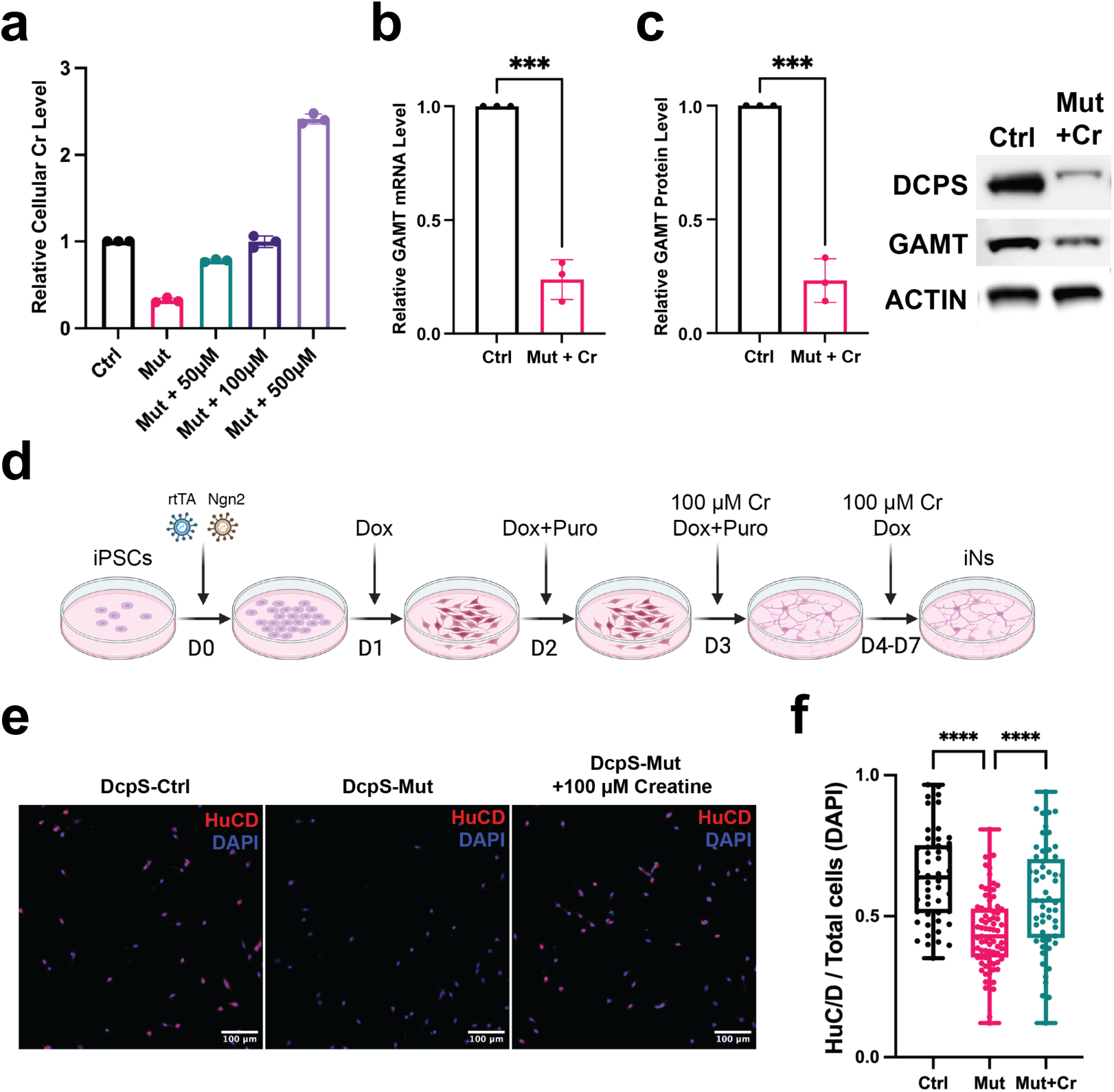
Creatine mediated reversal of DcpS mutant directed impairment of *in vitro* neurogenesis. **(a)** Levels denote intracellular Cr levels assessed by the Sigma Cr assay. **(b)** Cr supplementation does not impact GAMT mRNA levels. GAMT mRNA levels in DcpS- Mut cells complemented with 100µM Cr are shown normalized to GAPDH mRNA expression. Statistical analysis conducted on data from n = 3 biological replicates, each with three technical replicates, using an unpaired t-test. Significance represented by ***P < 0.001. **(C)** Bar graph of GAMT protein levels determined by western blot analysis in DcpS-Mut cells supplemented with 100µM Cr. Values are normalized to ACTIN protein levels. Statistical analysis performed on data from n = 3 biological replicates, with three technical replicates for each, using an unpaired t-test. Significance indicated by ***P < 0.001. **(d)** Schematic representation of DcpS-Mut human iPSC differentiation into induced neurons (iNs) and addition of Cr. Dox denotes doxycycline and Puro represents puromycin. **(e)** Immunostaining of DcpS human iNs at Day 7, depicting the expression of the neuronal marker HuC/D (in red) and nuclei stained with DAPI (in blue). **(f)** Box and Whisker plot illustrating the ratio of HuC/D-positive cells to DAPI-positive cells per image at Day 7. Imaging conducted at 20× magnification using the INCell Analyzer 6000 and analyzed with the INCarta™ software. Statistical analysis performed on data from DcpS-Ctrl (n = 50 images), DcpS-Mut (n = 77 images), and DcpS-Mut+Cr (n = 57 images) across three independent experiments. Significance determined by unpaired t-test, with ****P < 0.0001 denoting highly significant differences.

We next implemented the approach outlined in Figure 3D to differentiate iPSCs into iN cells in the presence or absence of Cr supplementation. Three days post transduction to initiate DcpS-Mut iPSCs differentiation, 100µM of Cr was supplemented into the culture medium and cells permitted to progress into iN cells. Monitoring neuronal differentiation was assessed through the expression of HuC/D, an RNA-binding protein specifically expressed in neurons and a marker for neuronal differentiation ^17,23,24^. Consistent with previous studies ^17^, immunofluorescent detection of HuC/D at post induction day 7 (D7) (Figure 3E) and high content analysis of microscopy images revealed a decrease in neuronal differentiation in DcpS mutant cells compared to control cells (Figure 3F). Interestingly, a significant increase in the number of mutant cells with elevated HuC/D levels was observed upon Cr supplementation, implying Cr supplementation mitigated the attenuated neurogenesis observed in iN cells derived from DcpS mutation-bearing iPSCs.

### Creatine Administration enhances Neurite Outgrowth in DcpS mutant iN cells

A second phenotypic characteristic of DcpS-Mut cells is impaired outgrowth of neuritic projections ^17^. Neuritogenesis was assessed by total immunofluorescence intensities of the neural-specific microtubule protein, β-III-tubulin (Tuj1) (Figure 4A). Quantification of aggregate neurite lengths showed a significant decrease in neuritic projection lengths in DcpS-Mut differentiated iN cells relative to DcpS-Ctrl cells (Figure 4B). Importantly, complementation with physiological levels of Cr substantially reversed the reduction (Figures 4A and 4B). Similarly, assessment of total Tuj1 intensity followed the same pattern and was reduced in the mutant cell and reversed upon supplementation with Cr (Figures 4A and 4C). Collectively, our findings suggest that Cr deficiency in DcpS mutant cells underlies the observed neurodevelopmental effects. Importantly, Cr supplementation could potentially reverse these effects, highlighting a therapeutic avenue for individuals with disrupted DcpS decapping activity.

**Figure 4:**
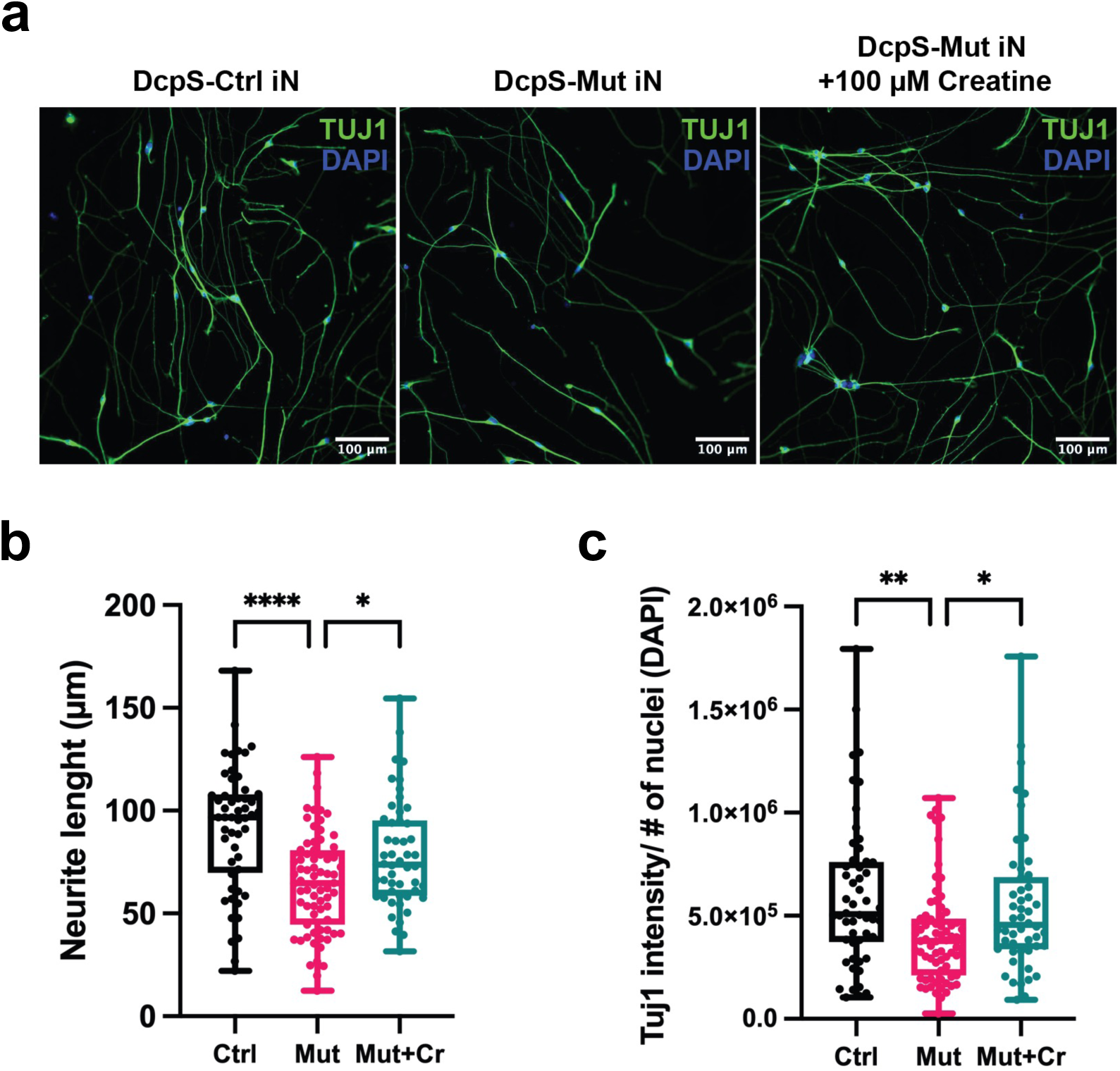
Creatine mediated reversal of DcpS mutant directed impairment of *in vitro* neuritogenesis. **(a)** Immunostaining of DcpS human iNs at Day 7, highlighting the expression of neuron-specific Class III β-tubulin (Tuj1) (in green) to label neurites. DAPI-positive nuclei are shown in blue **(b)** Box and Whisker plot illustrating the average neurite length (Tuj1 staining) per image for DcpS-Ctrl and DcpS-Mut Day 7 iNs. Imaging was performed at 20× magnification and analyzed using the INCarta™ image analysis software. Statistical analysis conducted on data from DcpS- Ctrl (n = 52 images), DcpS-Mut (n = 71 images), and DcpS-Mut+Cr (n = 50 images) across three independent experiments. Significance assessed by unpaired t-test, with ****P < 0.0001 and *P < 0.05. Statistical analysis was conducted using GraphPad Prism 9 software. **(c)** Box and Whisker plot representing the total Tuj1 intensity at Day 7 iNs, normalized to the number of nuclei in each image. Imaging and statistical analysis are as in panel B with **P < 0.01.

## Discussion

Our study provides significant insights into the molecular mechanisms underlying the neurodevelopmental disorders associated with the DcpS splice site mutation (c.636+1G>A). RNA sequencing revealed substantial alterations in the expression profiles of various transcripts in lymphocytes derived from patients carrying this mutation, notably downregulation of genes involved in the Cr biosynthesis pathway, such as GAMT, CKB and CKMT1B, and upregulation of those related to cell-cell signaling, including EPS8 and EPH4.

Metabolomics analysis further substantiated altered Cr synthesis pathway, showing marked reductions in intracellular Cr and Phospho-Cr levels with a concomitant elevation in GAA levels in DcpS mutant patient-derived cells that could be attributed to a disruption of GAMT activity. The significant downregulation of GAMT transcripts and protein in DcpS mutant cells indicated that the mutation hampers the conversion of GAA to Cr, leading to an accumulation of GAA and a decrease in the downstream metabolites, Cr and Phospho-Cr (Figure 2B).

Cr is an important cellular energy currency. In its phosphorylated form, Phosphor-Cr, is a critical energy storage source for the regeneration of ATP. Moreover, in the nervous system Cr can be neuroprotective ^25^ and can also function as a neurotransmitter ^26,27^ through modulation of GABAergic neurons ^28^. Consistent with the neural role for Cr, our findings demonstrate a critical dependence on Cr in the normal differentiation of DcpS-Mut iPSCs into iN cells. Most striking and important finding of our studies is the observation that the defects in neurogenesis and neuritogenesis observed in the DcpS-Mut iPSC differentiation into iNs are significantly reversed upon restoration of intracellular Cr to levels comparable to control cells. This is illustrated by the increased numbers of HuC/D-positive cells (Figures 3E, 3F) and enhanced neurite length (Figures 4) that are apparent following Cr supplementation. These results suggest DcpS-Mut cells harbor an underlying Cr deficiency, and this deficiency can be reversed. In particular, the increased number of HuC/D-positive cells and enhanced neurite lengths in Cr-supplemented cells underscore the efficacy of this intervention, suggesting that the neurodevelopmental deficits observed in DcpS mutant cells are due to impaired Cr biosynthesis. By bypassing the metabolic block caused by reduced GAMT levels, exogenous Cr mitigates the downstream physiological effects of the mutation.

Interestingly, a spectrum of disorders collectively termed Creatine Deficiency Syndromes (CDS) encompass a wide range of metabolic disorders characterized by impaired Cr synthesis or transport, crucial for energy metabolism in the brain and muscles ^29,30^. They manifest as defects in either of two enzymes that are necessary for Cr production, L-arginine-glycine amidinotransferase which generates GAA, or GAMT which converts the GAA precursor into Cr (Figure 2B). Moreover, in addition to reduced Cr levels, deficiencies in GAMT results in the accumulation of neurotoxic GAA, significantly impacting neurological function ^31^. A defect in a third gene class that constitutes CDS is *SLC6A8*, a membrane bound Cr transporter necessary for Cr uptake into organs. Importantly, two common neurological defects in CDS individuals are cognitive disability and delays in speech ^22^, both of which are evident in the DcpS mutant individuals ^13^.

Our studies indicated that the observed reduction of GAMT expression, elevated GAA and reduction in Cr metabolites in the DcpS-Mut cells are consistent with CDS. We propose individuals harboring impaired DcpS decapping can be characterized within the class of Creatine Deficiency Syndromes and their Cr deficiency is likely a contributor of neurodevelopmental disorder. It should be noted that although addition of Cr significantly reversed the diminished neurogenesis and neuritogenesis, it did not completely restore it, indicating that additional factors are also likely involved. For example, reductions of NAT8L ((N-acetyltransferase-8-like protein Nat8L) also contribute to curtailed neurogenesis ^32^, indicating DcpS may be a broader coordinator of neurogenesis. Nevertheless, these findings suggest that similar to other GAMT defective conditions that benefit from bypassing the metabolic block through therapeutic supplementation with Cr ^31^, a similar strategy may be beneficial for individuals with DcpS decapping mutations likely most beneficial prior to irreversible neurologic damage.

In conclusion, our study elucidates the underpinnings of the neurodevelopmental disorder associated with the DcpS splice site mutation and identifies the Cr biosynthetic pathway as at least one critical contributor to the underlying neurodevelopmental defect. These finding raise the intriguing possibility of Cr as a potential therapeutic modality to ameliorate the neurological deficits associated with this mutation. These studies also raise important questions for future inquiry in delineating the molecular mechanism of how a disruption of DcpS decapping leads to a decrease in GAMT mRNA levels which appears as the major catalyst for establishing the phenotype.

## Acknowledgments

We are grateful to Eric Chiles and Xiaoyang Su for their assistance with the mass spectrometry analysis. This work was supported by National Institutes of Health (NIH) grant GM149262 (M.K.).

## Data availability

All unique materials and reagents generated in this study are available from the corresponding author with a completed material transfer agreement. The sequencing data is deposited at NIH GEO (GSE274722). Any additional information required to reanalyze the data reported in this paper is available from the corresponding author upon request.

## Methods

### Neuronal Differentiation of iPSCs

Lymphoblast cells from individuals IV-3, harboring a homozygous DcpS mutation (DcpS-Mut), and the DcpS mutant heterozygote individual III-3 (DcpS-Cntl), were reported in Ahmed et al. ^13^, and their derived iPSCs are reported in Salamon et al. ^17^. Briefly, the iPSCs were generated by RUCDR Infinite Biologics® using Sendai viral vectors (CytoTune™, Thermo Fisher Scientific), as previously described (Moore et al. 2012). The iPSCs were maintained in mTeSR™ Plus hPSC medium (STEMCELL Technologies). Cells were passaged using either ReLeSR™ passaging reagent (STEMCELL Technologies) or Accutase (STEMCELL Technologies) and cultured onto Matrigel-coated six-well plates.

Induced cortical neurons (iNs) were generated directly from human iPSCs using an inducible lentiviral system, as described ^17^. Briefly, iPSCs were plated in the presence of the Y-compound (5 μM) and transduced with lentiviruses encoding rtTA (FUW-M2rtTA) and NGN2 (Tet-O-Ngn2- puro). NGN2 expression was induced the following day by adding doxycycline (2 μg/mL) to neurobasal culture medium containing B27 supplement (Gibco). Infected cells were selected for 48 hours using puromycin (2 μg/mL) and subsequently cultured for an additional 5–7 days in B27- and doxycycline-containing neurobasal medium. All experiments with the cell lines were carried out in accordance with relevant guidelines and experimental approved protocols by the Rutgers University Institutional Review Board (IRB #15-787M) with informed consent of subject guardians.

### Immunofluorescence and Microscopy Analysis

Undifferentiated iNs (day 0) and those differentiated for 7 days were processed as follows: cells were washed with cold 1× PBS, fixed with 4% paraformaldehyde, permeabilized with 0.1% Triton X-100, and incubated in a blocking buffer containing 10% normal goat serum. Cells were then exposed to primary antibodies diluted in a PBS buffer containing 10% normal goat serum for 1 hour at room temperature. Primary antibodies included HuC/D (Thermo Fisher, #A-21271; 1:250) and TUJ1 (BioLegend, #801202; 1:250), incubated overnight at 4°C. After washing with 1× PBS, cells were incubated with secondary antibodies for 1 hour, followed by another round of washing with 1× PBS and staining with DAPI (Thermo Fisher; 1:1000). Secondary antibodies included goat anti-mouse IgG conjugated to AlexaFluor-647 (Thermo Fisher, #A-21235) and goat anti- rabbit IgG conjugated to AlexaFluor-488 (Thermo Fisher, #A-11008).

To assess neuronal differentiation, immunostained cells for HuC/D or Tuj1 were imaged using the INCell Analyzer 6000 (GE Healthcare) at 20x magnification. Images with a comparable number of nuclei per image, obtained from three independent experiments, were analyzed using the INCarta™ image analysis software (GE Healthcare). Statistical analysis and graphical representation were performed using GraphPad Prism (version 9) software. An unpaired t-test was used for comparing subject groups, given the normal distribution of values.

### Western Blotting

Cells were lysed in a phosphate-buffered saline solution containing 0.1% Tween 20 (PBST) and protease inhibitors (Sigma-Aldrich, #11873580001), followed by sonication. Equivalent protein quantities from various samples were separated using Bolt™ 4-12% Bis-Tris Plus Gels (Thermo Fisher Scientific, #NW04120BOX) and transferred onto nitrocellulose membranes (Bio-Rad, #1620115). Membranes were exposed to primary antibodies (hDCPS anti-rabbit, GAMT Proteintech 10880-1-AP) in PBST with 5% BSA (Sigma-Aldrich), followed by secondary antibodies tagged with horseradish peroxidase. Protein visualization was achieved using the ECL Prime Western blotting detection reagent (Cytiva, #RPN2232).

### RNA-Seq Analysis

Total cellular RNA was harvested with TRIzol Reagent (Thermo Fisher Scientific) and treated with RNase-free DNase (Promega) following manufacturers’ protocols. Ten micrograms of total RNA were flash-frozen in liquid nitrogen and sequenced at GENEWIZ (Azenta Life Sciences). Results (fastq files) were trimmed and de-duplicated using fastp (v0.23.4)^33^, aligned with reference genome (GRCh38) using HISAT2 (v2.2.0)^34^, and gene counts extracted from bam files using the featureCounts function from the Rsubread package (v2.16.1)^35^. Counts were scaled, normalized and fit to a model using DESeq2 (v1.42.1)^36^. Differentially expressed genes were identified by Wald testing, FDR-adjusted p < 0.05 and log2 fold change > 1. Data are available from NIH GEO (GSE274722).

### Reverse-Transcription (RT)-qPCR Analysis of Gene Expression

RNA was extracted using TRIzol™ (Thermo Fisher Scientific) followed by isopropanol precipitation. Extracted RNA was treated with DNase I (Promega) and used for reverse transcription (Promega RT kit) to generate cDNA. Gene expression analysis was performed using SYBR Green dye on an Applied Biosystems QuantStudio 3 qPCR machine (Thermo Fisher Scientific). GAPDH mRNA was used as a housekeeping gene for normalization. The relative gene expression levels were calculated using the delta-delta Ct (ΔΔCt) method. Briefly, the Ct values of the target genes were normalized to the Ct values of GAPDH (ΔCt), and the ΔCt values were then compared to a control sample to obtain the ΔΔCt values. The fold change in gene expression was determined using the 2^(-ΔΔCt) method.

### Cell culture and Metabolite Extraction

Patient-derived lymphoblasts were cultured in RPMI 1640 medium. For metabolite extraction, cells from 10 cm plates were harvested by centrifugation at 300g for 5 minutes, followed by washing with 1xPBS. Metabolic activity was quenched using an extraction buffer composed of 40:40:20 (v/v) Acetonitrile:Methanol:Water (Thermo Fisher Scientific, Waltham, MA) with 0.1M Formic acid (40 mL of extraction solvent per 1 mL of packed cell volume) and Eppendorf tubes were placed on dry ice for 10 minutes. Subsequently, the extraction solution was neutralized with 15% (m/v) Ammonium Bicarbonate (70 µL for 800 µL of extraction buffer). The tubes were then centrifuged in a benchtop microcentrifuge at maximum speed for 30 minutes at 4°C. The resulting supernatant was collected and transferred to LC-MS vials for analysis.

### Ultra-High Performance Liquid Chromatography conditions

The HILIC separation was performed on a Vanquish Horizon UHPLC system (Thermo Fisher Scientific, Waltham, MA) with XBridge BEH Amide column (150 mm × 2.1 mm, 2.5 μm particle size, Waters, Milford, MA) using a gradient of solvent A (95%:5% H2O:acetonitrile with 20 mM acetic acid, 40 mM ammonium hydroxide, pH 9.4), and solvent B (20%:80% H2O:acetonitrile with 20 mM acetic acid, 40 mM ammonium hydroxide, pH 9.4). The gradient was 0 min, 100% B; 3 min, 100% B; 3.2 min, 90% B; 6.2 min, 90% B; 6.5 min, 80% B; 10.5 min, 80% B; 10.7 min, 70% B; 13.5 min, 70% B; 13.7 min, 45% B; 16 min, 45% B; 16.5 min, 100% B and 22 min, 100% B ^37^ at a flow rate of 300μl/min. Injection volume was 5 μL and column temperature was 25 °C. The autosampler temperature was set to 4°C and the injection volume was 5µL.

### Full scan mass spectrometry

The full scan mass spectrometry analysis was performed on a Thermo Q Exactive PLUS with a HESI source which was set to a spray voltage of -2.7kV under negative mode and 3.5kV under positive mode. The sheath, auxiliary, and sweep gas flow rates of 40, 10, and 2 (arbitrary unit) respectively. The capillary temperature was set to 300°C and aux gas heater was 360°C. The S-lens RF level was 45. The m/z scan range was set to 72 to 1000m/z under both positive and negative ionization mode. The AGC target was set to 3e6 and the maximum IT was 200ms. The resolution was set to 70,000.

### Mass spectrometry data analysis and data quality

The full scan data was processed with a targeted data pipeline using MAVEN software package ^38^. The compound identification was assessed using accurate mass and retention time match to the metabolite standards from the in- house library. Prior to running the samples, the LC-MS system was evaluated for performance readiness by running a commercially available standard mixture and an in-house standard mixture to assess the mass accuracy, signal intensities, and retention time consistency. All known metabolites in the mixture are detected within 5ppm mass accuracy. Method blank samples matching the composition of the extraction solvent are used in every sample batch to assess background signals and ensure there isn’t carryover from one run to the next. In addition, the sample queue was randomized with respect to sample treatment to eliminate the potential for batch effects.

### Quantification of Intracellular Cr Levels

Lymphoblast cells were seeded at a density of 2 x10^6^ cells per 10 cm plate and exposed to varying concentrations of Cr (0, 0.1, 0.5, 1, 2, 3, and 5 mM) for 24 hours. Following incubation, cells were washed thoroughly with phosphate-buffered saline (PBS) five times to remove residual Cr. The cells were then harvested by centrifugation and homogenized in 400 µL of assay buffer. To prepare the samples for Cr quantification, the homogenates were subjected to centrifugation at 13,000 g for 10 minutes at 4°C using an Amicon 10 kDa molecular weight cutoff (MWCO) spin filter. The Cr concentration in the filtered samples was determined using the Cr Assay Kit (Sigma-Aldrich, Cat. #MAK079-1KT).

## References

1. Grudzien-Nogalska, E., Jiao, X., Song, M.G., Hart, R.P. & Kiledjian, M. Nudt3 is an mRNA decapping enzyme that modulates cell migration. RNA 22, 773–81 (2016).

2. Husain, R.A. et al. Biallelic NUDT2 variants defective in mRNA decapping cause a neurodevelopmental disease. Brain 147, 1197–1205 (2024).

3. Lykke-Andersen, J. Identification of a human decapping complex associated with hUpf proteins in nonsense-mediated decay. Mol Cell Biol 22, 8114–21 (2002).

4. Song, M.G., Li, Y. & Kiledjian, M. Multiple mRNA decapping enzymes in mammalian cells. Mol Cell 40, 423–32 (2010).

5. Wang, Z., Jiao, X., Carr-Schmid, A. & Kiledjian, M. The hDcp2 protein is a mammalian mRNA decapping enzyme. Proc Natl Acad Sci U S A 99, 12663–8. (2002).

6. Song, M.G., Bail, S. & Kiledjian, M. Multiple Nudix family proteins possess mRNA decapping activity. RNA 19, 390–9 (2013).

7. Doamekpor, S.K., Sharma, S., Kiledjian, M. & Tong, L. Recent insights into noncanonical 5’ capping and decapping of RNA. J Biol Chem 298, 102171 (2022).

8. Liu, H., Rodgers, N.D., Jiao, X. & Kiledjian, M. The scavenger mRNA decapping enzyme DcpS is a member of the HIT family of pyrophosphatases. EMBO J 21, 4699–708 (2002).

9. Milac, A.L., Bojarska, E. & Wypijewska del Nogal, A. Decapping Scavenger (DcpS) enzyme: advances in its structure, activity and roles in the cap-dependent mRNA metabolism. Biochim Biophys Acta 1839, 452–62 (2014).

10. Wang, Z. & Kiledjian, M. Functional Link between the Mammalian Exosome and mRNA Decapping. Cell 107, 751–762. (2001).

11. Wulf, M.G. et al. The yeast scavenger decapping enzyme DcpS and its application for in vitro RNA recapping. Sci Rep 9, 8594 (2019).

12. Prashad, S. & Gopal, P.P. RNA-binding proteins in neurological development and disease. RNA Biol 18, 972–987 (2021).

13. Ahmed, I. et al. Mutations in DCPS and EDC3 in autosomal recessive intellectual disability indicate a crucial role for mRNA decapping in neurodevelopment. Hum Mol Genet 24, 3172–80 (2015).

14. Alesi, V. et al. An additional patient with a homozygous mutation in DCPS contributes to the delination of Al-Raqad syndrome. Am J Med Genet A 176, 2781–2786 (2018).

15. Masoudi, M. et al. Leukoencephalopathy in Al-Raqad syndrome: Expanding the clinical and neuroimaging features caused by a biallelic novel missense variant in DCPS. Am J Med Genet A 182, 2391–2398 (2020).

16. Ng, C.K. et al. Loss of the scavenger mRNA decapping enzyme DCPS causes syndromic intellectual disability with neuromuscular defects. Hum Mol Genet 24, 3163–71 (2015).

17. Salamon, I. et al. mRNA-Decapping Associated DcpS Enzyme Controls Critical Steps of Neuronal Development. Cereb Cortex 32, 1494–1507 (2022).

18. Wallimann, T., Tokarska-Schlattner, M. & Schlattner, U. The creatine kinase system and pleiotropic effects of creatine. Amino Acids 40, 1271–96 (2011).

19. Wallimann, T., Wyss, M., Brdiczka, D., Nicolay, K. & Eppenberger, H.M. Intracellular compartmentation, structure and function of creatine kinase isoenzymes in tissues with high and fluctuating energy demands: the ’phosphocreatine circuit’ for cellular energy homeostasis. Biochem J 281 **( Pt** **1**), 21–40 (1992).

20. Beard, E. & Braissant, O. Synthesis and transport of creatine in the CNS: importance for cerebral functions. J Neurochem 115, 297–313 (2010).

21. Forbes, S.C. et al. Effects of Creatine Supplementation on Brain Function and Health. Nutrients 14(2022).

22. Stockler, S., Schutz, P.W. & Salomons, G.S. Cerebral creatine deficiency syndromes: clinical aspects, treatment and pathophysiology. Subcell Biochem 46, 149–66 (2007).

23. Chung, S., Jiang, L., Cheng, S. & Furneaux, H. Purification and properties of HuD, a neuronal RNA-binding protein. J Biol Chem 271, 11518–24 (1996).

24. Zucco, A.J. et al. Neural progenitors derived from Tuberous Sclerosis Complex patients exhibit attenuated PI3K/AKT signaling and delayed neuronal differentiation. Mol Cell Neurosci 92, 149–163 (2018).

25. Bahari, Z., Jangravi, Z., Hatef, B., Valipour, H. & Meftahi, G.H. Creatine supplementation protects spatial memory and long-term potentiation against chronic restraint stress. Behav Pharmacol 34, 330–339 (2023).

26. Libell, J.L. et al. Guanidinoacetate N-methyltransferase deficiency: Case report and brief review of the literature. Radiol Case Rep 18, 4331–4337 (2023).

27. Meftahi, G.H., Hatef, B. & Pirzad Jahromi, G. Creatine Activity as a Neuromodulator in the Central Nervous System. Arch Razi Inst 78, 1169–1175 (2023).

28. Schulze, A., Tran, C., Levandovskiy, V., Patel, V. & Cortez, M.A. Systemic availability of guanidinoacetate affects GABAA receptor function and seizure threshold in GAMT deficient mice. Amino Acids 48, 2041–7 (2016).

29. Goldstein, J. et al. ClinGen variant curation expert panel recommendations for classification of variants in GAMT, GATM and SLC6A8 for cerebral creatine deficiency syndromes. Mol Genet Metab 142, 108362 (2024).

30. Schulze, A. Creatine deficiency syndromes. Handb Clin Neurol 113, 1837–43 (2013).

31. Fernandes-Pires, G. & Braissant, O. Current and potential new treatment strategies for creatine deficiency syndromes. Mol Genet Metab 135, 15–26 (2022).

32. Wulaer, B. et al. Shati/Nat8l deficiency disrupts adult neurogenesis and causes attentional impairment through dopaminergic neuronal dysfunction in the dentate gyrus. J Neurochem 157, 642–655 (2021).

33. 33. Chen, S. Ultrafast one-pass FASTQ data preprocessing, quality control, and deduplication using fastp. Imeta 2, e107 (2023).

34. Kim, D., Langmead, B. & Salzberg, S.L. HISAT: a fast spliced aligner with low memory requirements. Nat Methods 12, 357–60 (2015).

35. Liao, Y., Smyth, G.K. & Shi, W. The R package Rsubread is easier, faster, cheaper and better for alignment and quantification of RNA sequencing reads. Nucleic Acids Res 47, e47 (2019).

36. Love, M.I., Huber, W. & Anders, S. Moderated estimation of fold change and dispersion for RNA-seq data with DESeq2. Genome Biol 15, 550 (2014).

37. Su, X. et al. In-Source CID Ramping and Covariant Ion Analysis of Hydrophilic Interaction Chromatography Metabolomics. Anal Chem 92, 4829–4837 (2020).

38. Melamud, E., Vastag, L. & Rabinowitz, J.D. Metabolomic analysis and visualization engine for LC-MS data. Anal Chem 82, 9818–26 (2010).

## Methods References

1. Ahmed, I., Buchert, R., Zhou, M., Jiao, X., Mittal, K., Sheikh, T.I., Scheller, U., Vasli, N., Rafiq, M.A., Brohi, M.Q., et al. (2015). Mutations in DCPS and EDC3 in autosomal recessive intellectual disability indicate a crucial role for mRNA decapping in neurodevelopment. Hum Mol Genet 24, 3172–3180. 10.1093/hmg/ddv069.

2. Salamon, I., Palsule, G., Luo, X., Roque, A., Tucai, S., Khosla, I., Volk, N., Liu, W., Cui, H., Pozzo, V.D., et al. (2022). mRNA-Decapping Associated DcpS Enzyme Controls Critical Steps of Neuronal Development. Cereb Cortex 32, 1494–1507. 10.1093/cercor/bhab302.

3. Chen, S. (2023). Ultrafast one-pass FASTQ data preprocessing, quality control, and deduplication using fastp. Imeta 2, e107. 10.1002/imt2.107.

4. Kim, D., Langmead, B., and Salzberg, S.L. (2015). HISAT: a fast spliced aligner with low memory requirements. Nat Methods 12, 357–360. 10.1038/nmeth.3317.

5. Liao, Y., Smyth, G.K., and Shi, W. (2019). The R package Rsubread is easier, faster, cheaper and better for alignment and quantification of RNA sequencing reads. Nucleic Acids Res 47, e47. 10.1093/nar/gkz114.

6. Love, M.I., Huber, W., and Anders, S. (2014). Moderated estimation of fold change and dispersion for RNA-seq data with DESeq2. Genome biology 15, 550. 10.1186/s13059-014- 0550-8.

7. Su, X., Chiles, E., Maimouni, S., Wondisford, F.E., Zong, W.X., and Song, C. (2020). In-Source CID Ramping and Covariant Ion Analysis of Hydrophilic Interaction Chromatography Metabolomics. Anal Chem 92, 4829–4837. 10.1021/acs.analchem.9b04181.

8. Melamud, E., Vastag, L., and Rabinowitz, J.D. (2010). Metabolomic analysis and visualization engine for LC-MS data. Anal Chem 82, 9818–9826. 10.1021/ac1021166.

